# Interspecific territoriality has facilitated recent increases in the breeding habitat overlap of North American passerines

**DOI:** 10.1101/2022.08.23.504954

**Authors:** D.A. Nesbit, M.C. Cowen, G.F. Grether, J.P. Drury

## Abstract

As species’ ranges shift in response to human-induced global changes, species interactions are expected to play a large role in shaping the resultant range dynamics and, subsequently, the composition of modified species assemblages. Most research on the impact of species interactions on range dynamics focuses on the effects of trophic interactions and exploitative competition for resources, but an emerging body of work shows that interspecific competition for territories and mates also affects species range shifts. As such, it is paramount to build a strong understanding of how these forms of behavioural interference between species impact landscape-scale patterns. Here, we examine recent (1997-2019) range dynamics of North American passerines to test the hypothesis that behavioural interference impacts the ease with which species move across landscapes. Over this 22-year period, we found that fine-scale spatial overlap between species (syntopy) increased more for species pairs that engage in interspecific territoriality than for those that do not. We found no evidence, however, for an effect of reproductive interference (hybridisation) on syntopy, and no effect of either type of interference on range-wide overlap (sympatry). Examining the net effects of species interactions on continent-scale range shifts may require species occurrence data spanning longer time periods than are currently available for North American passerines, but our results show that interspecific territoriality has had an overall stabilising influence on species coexistence over the past two decades.

## INTRODUCTION

Species ranges are changing as a result of climate change, land-use change, and introduction to non-native areas (Thomas and Lennon 1999, La Sorte and Boecklen 2005a, Hitch and Leberg 2007, Zuckerberg et al. 2009, Brommer et al. 2012, Elmhagen et al. 2015, Dyer et al. 2017). Such movement across landscapes is expected to be impacted (either positively or negatively) by interactions between currently coexisting taxa (e.g., Svenning et al. 2014). Similarly, shifts into new areas may give rise to novel species interactions, which may in turn determine the likelihood that a range-shifting lineage successfully becomes established (HilleRisLambers et al. 2013, Svenning et al. 2014, Sirén and Morelli 2020). The impact of such species interactions on range dynamics has been investigated extensively for interactions between trophic levels (e.g., predator-prey or plant-pollinator interactions; Wisz et al. 2013, Svenning et al. 2014) and within trophic levels (e.g., exploitative competition or facilitation; Connor and Bowers 1987, Heikkinen et al. 2007, Wisz et al. 2013, Svenning et al. 2014, Ortego & Knowles 2020, Novella-Fernandez et al. 2021). Yet, in addition to competition for resources, many animals engage in behavioural interference, a widespread phenomenon that encompasses interspecific interactions such as interference competition (e.g. interspecific territoriality) and reproductive interference (e.g. hybridisation) (Connor and Bowers 1987, Gross and Price 2000, Gotelli et al. 2010, Krosby and Rohwer 2010, McQuillan and Rice 2015, Grether et al. 2017).

An emerging body of research suggests that behavioural interference between species can influence the outcome of range shifts (Gross and Price 2000, Duckworth and Badyaev 2007, Jankowski et al. 2010, McQuillan and Rice 2015). The hypothesised impacts of such interference largely depend on the fitness costs and benefits of engaging in territorial or reproductive interactions with heterospecifics. On the one hand, when behavioural interference incurs net fitness costs for one or more interacting species, these costs diminish population growth and lead to an increased risk of local extinction (i.e., sexual or competitive exclusion; Kuno 1992, Liou and Price 1994, Amarasekare 2002, Hochkirch et al. 2007, Gröning and Hochkirch 2008, Pfennig and Pfennig 2012, Kishi and Nakazawa 2013, Legault et al. 2020). On the other hand, if behavioural interference instead diminishes interspecific resource competition by reducing spatial overlap, such interference may enable ecologically similar species pairs to coexist (Case and Gilpin 1974, Zhang and Hanski 1998, Mikami and Kawata 2004, Grether et al. 2013, Kishi and Nakazawa 2013, Ruokolainen and Hanski 2016, Gómez-Llano et al. 2021, Grether and Okamoto 2022).

Empirical research confirms that behavioural interference can lead to competitive or reproductive exclusion at a variety of scales (Robinson and Terborgh 1995, Gross and Price 2000, Duckworth and Badyaev 2007, Jankowski et al. 2010, Grether et al. 2013, Pasch et al. 2013, Grether et al. 2017, Freeman et al. 2019, 2022). At local scales, behavioural interference can lead to the exclusion of species from particular habitat patches so that species pairs coexist in sympatry but not syntopy (Robinson and Terborgh 1995, Vallin et al. 2012, Rybinski et al. 2016, Reif et al. 2018). For instance, collared flycatchers (*Ficedula albicollis*) that have colonised the Swedish island of Öland in the last 50 years are competitively dominant over pied flycatchers (*Ficeldula hypoleuca*) and confine them to lower quality coniferous woodlots through both interspecific territoriality and reproductive interference via costly hybridisation (Vallin and Qvarnström 2011, Vallin et al. 2012, Rybinski et al. 2016). At larger scales, more extensive exclusion occurs at geographical range boundaries when behavioural interference precludes coexistence in sympatry (Gross and Price 2000, Duckworth and Badyaev 2007, Jankowski et al. 2010, Krosby and Rohwer 2010, McQuillan and Rice 2015, Freeman et al. 2016, Freeman and Montgomery 2016, Legault et al. 2020). For instance, several studies of montane species pairs occupying abutting altitudinal ranges suggest that interspecific territoriality, rather than abiotic factors and differing habitat requirements, is the key factor preventing range overlap at elevational boundaries (Jankowski et al. 2010, Freeman and Montgomery 2016, Freeman et al. 2016, 2019, 2022, Boyce and Martin 2019). Behavioral interference has also been implicated as the primary factor limiting species ranges along latitudinal and habitat gradients (Gross and Price 2000, Duckworth and Badyaev 2007, Krosby and Rohwer 2010, McQuillan and Rice 2015, Martin and Bonier 2018).

The notion that behavioural interference might facilitate coexistence has received less empirical attention, but nevertheless, some studies directly support this idea (Marvin 1998, Ovadia and Zu Dohna 2003, Ziv and Kotler 2003, Levy et al. 2011, Pasch et al. 2013). Several other studies demonstrate stable coexistence between ecologically similar species that are interspecifically territorial (Rohwer 1973, Reed 1982, Jankowski et al. 2012, Drury et al. 2015, 2019, Freeman 2016, Boyce and Martin 2019, Grether et al. 2020), which is consistent with the hypothesis that interspecific territoriality can enable resource partitioning, thereby enabling coexistence. For instance, chaffinches (*Fringilla coelebs*) and great tits (*Parus major*) on the mainland of Scotland occupy similar niches, exist in overlapping territories, and do not respond aggressively to heterospecific playback (Reed 1982). However, on adjacent islands they defend non-overlapping interspecific territories and respond aggressively to heterospecific playback (Reed 1982). This interspecific territoriality may allow them to coexist on relatively resource-poor islands without the competitive exclusion of either species. Further compelling evidence that behavioural interference can lead to stable coexistence comes from examples of convergent character displacement acting on territorial signals (Cody 1969, Tobias and Seddon 2009, Kirschel et al. 2019, Miller et al. 2019). Eastern (*Sturnella magna*) and western meadowlarks (*Sturnella neglecta*), for example, have converged in plumage patterning and colouration where they co-occur and defend non-overlapping interspecific territories (Rohwer 1973).

Consistent with theoretical work showing that interspecific territoriality can stabilise coexistence between ecologically similar species (Grether and Okamoto 2022), recent comparative analyses on North American passerines have shown that interspecific territoriality is positively associated with resource use and fine-scale habitat overlap (Cowen et al. 2020, Drury et al. 2020). Here, we test additional predictions that follow from the hypothesis that interspecific territoriality can stabilise coexistence. First, in taxa with shifting or expanding ranges, the extent of range overlap (sympatry) should increase more over time between interspecifically territorial species compared to non-interspecifically territorial species. Second, regardless of whether species ranges are changing, the magnitude of fine-scale habitat overlap (syntopy) within the areas of range overlap should increase more (or decrease less) over time between interspecifically territorial species compared to non-interspecifically territorial species. The second prediction does not only apply to taxa with expanding ranges, but it does assume that dispersal is a regular occurrence and that some suitable habitats within the species’ ranges remain unoccupied.

For hybridizing species, frequency-dependent interactions can generate Allee effects that might prevent one species from expanding into other species’ range (Kishi et al. 2009, Kyogoku and Nishida 2012, Kishi and Nakazawa 2013, Bargielowski and Lounibos 2016, Noriyuki and Osawa 2016, Whitton et al. 2017). In areas where the species’ ranges do overlap, however, selection against interspecific mating could cause fine-scale habitat partitioning (Gröning et al. 2007, Gómez-Llano et al. 2021), which in turn could reduce interspecific exploitative competition for resources and facilitate further increases in range overlap. Thus, it is difficult to predict whether hybridization would be positively or negatively associated with temporal changes in sympatry, but it seems robust to predict that hybridization should not be associated with temporal increases in syntopy.

To test the above predictions in North American passerines, we combined data on interspecific territoriality and hybridization (Drury et al. 2020) with data on recent range dynamics (1997-2019) (Sauer et al. 2020).

## METHODS

### Syntopy & sympatry

We calculated several indices of temporal change in interspecific spatial overlap using data from the North American Breeding Bird Survey (BBS) (Sauer et al. 2020) (Fig. 1). The BBS has been running since 1966 and is conducted by trained observers carrying out roadside point counts at 50 stops along each 39.4km route. As of 2019, 5756 routes had been surveyed (Supporting information Fig. S1), although not all routes had been surveyed annually (Sauer et al. 2020). Because data for each individual stop (i.e., the ‘50-stop’ dataset) is only available consistently for surveys conducted after 1996, and because the number of routes surveyed each year (i.e., the ‘10-stop’ dataset) plateaued around 1995 (Supporting information Fig. S2), for the sake of comparability, we focused our analyses on the years 1997-2019. Our analyses cover the trends and dynamics of species ranges within the BBS area; however, the ranges of some species extend beyond these limits (Supporting information Fig. S1).

**Figure 1.**
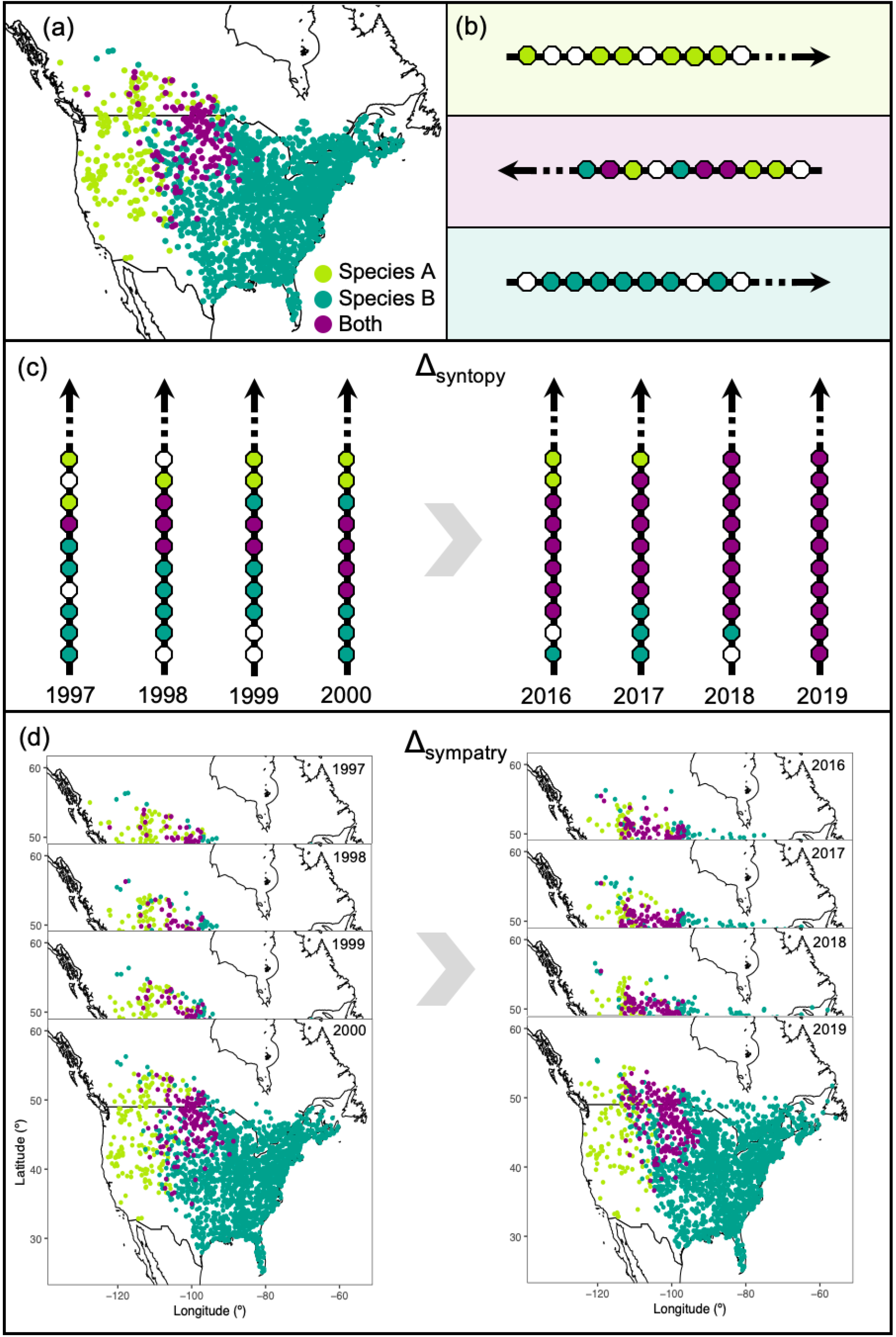
Measures of coocurrence using North American Breeding Bird Survey (BBS) data. (a) Each dot on the map represents a BBS route where only species A was present (green), where only species B was present (blue) or where both species were present (purple). (b) Each arrow represents one fifth of a hypothetical BBS route and each hexagon on the route represents a stop where only species A (green), only species B (blue), both species (purple) or neither species (white) were present. The purple box represents routes where species A and B co-occur in sympatry and syntopy while the green and blue boxes represent routes where the species are allopatric. (c) Syntopy was calculated as the number of stops shared by both species divided by the number of stops with only one present. This value for each route was then averaged across all routes at which both species were present within each year. Δ_syntopy_ was calculated by subtracting mean syntopy between 1997-2000 from mean syntopy 2016-2019. (d) Sympatry was calculated as the number of routes shared by both species divided by the number of routes occupied by the species occurring at the fewest and Δ_sympatry_ was calculated by subtracting mean sympatry 1997-2000 from mean sympatry 2016-2019.

From the 50-stop BBS data, we calculated a BBS-derived index of syntopy, or cooccurrence in the same breeding habitat within the zone of sympatry (Rivas 1964), as the number of stops shared by both species divided by the number of stops with only one present (Fig. 1b). We calculated this value for each route and averaged values across all routes at which both species were present within each year. Then, we calculated the temporal change in syntopy (Δ_syntopy_) for each species pair by subtracting the mean annual syntopy value between 1997-2000 from the mean annual syntopy between 2016-2019 (Fig. 1c).

Using the 10-stop BBS data, we calculated an index of sympatry (range overlap) as the number of routes with both species divided by the number of routes with the species that occurred on the fewest routes (similar to the Szymkiewicz-Simpson index used on range maps, e.g. Pigot et al. 2016) in each year (Fig. 1a). As above, we then calculated the temporal change in sympatry (Δ_sympatry_) by subtracting the mean of 1997-2000 sympatry values from the mean of 2016-2019 values (Fig. 1d). All BBS routes that were run in a given year were included in the syntopy and sympatry calculations.

### Interspecific behavioural interference

We used a database of interspecific territorial behaviour in North American passerines compiled by Drury et al. (2020). Briefly, species pairs in this database were considered to be interspecifically territorial if there were multiple reported territorial interactions between them in the literature. The database also includes a list of species pairs that were considered not to be interspecifically territorial based on census data from regions and time periods where observations of interspecific territorial behaviour were made (Losin et al. 2016, Drury et al. 2020). Hybridisation classifications were also taken from Drury et al. (2020) and did not include captive hybridisation or unsubstantiated reports of hybridisation. In total, we included 1602 out of 1618 pairs identified by Drury et al. (2020), as 16 pairs were not found on any of the same routes in both time periods. Of the 1602 pairs used in analyses, 74 pairs were interspecifically territorial, 1528 were not, 68 pairs hybridised whereas 1534 did not, and 27 pairs were both interspecifically territorial and hybridising.

### Indices of niche overlap

Foraging guild classifications, the square root of body mass, and the square root of bill length difference were also taken from Drury et al. (2020) and included as fixed effects as measures of niche overlap. Foraging guild components were classified according to De Graaf et al. (1985) and included the main dietary component, foraging technique, and foraging substrate. The proportion of overlap across foraging axes for each pair was calculated as a measure of foraging niche overlap. Species pairs that overlap in foraging guild are likely to occupy similar niches and therefore compete with one another (De Graaf et al. 1985), which may in turn influence coexistence (Gause 1934). Similarly, species with a similar body masses and bill lengths are also more likely to occupy similar niches (Pigot et al. 2020). Whether both members of a pair were secondary cavity nesters was also included as a fixed effect, as these species may compete for nest sites as well as food resources (Brawn and Balda 1988). Whether species in a pair share the same habitat type was included as a fixed effect as another indicator of potential niche overlap and coexistence. Habitat classifications were based on scores from Drury et al. (2020) and refer to the preferred habitat of each species. Categories 1-3 equate to open, semi-dense and dense habitats respectively. To test whether the type of breeding territory a species defends impacts changes in range overlap (Freeman et al. 2019), we included an index indicating whether both members of a pair defended the same intraspecific territory type (using a categorical index ranging from 1 [not territorial] to 5 [defending a ‘multipurpose’ territory in which both foraging and nesting occurs]; Drury et al. 2020). Patristic distance, the time separating members of a pair on the phylogeny, was included in models to control for the time available for evolutionary divergence between taxa (Tobias et al. 2014).

### Range expansion and contraction

We controlled for changes in range size, which may also impact changes in sympatry and syntopy, by generating indices of range expansion and range contraction. Specifically, if a species was present at a route between 1997 and 2000 but not present between 2016 and 2019, it was considered to have been extirpated from the route. Conversely, if a species was absent between 1997 and 2000 but present between 2016 and 2019, it was considered to have colonised the route. For each species, the change in range size was derived as the number of routes colonised between 1997-2000 and 2016-2019 minus the number of routes at which the species was extirpated (Supporting information Fig. S3). If this value was positive, a species was considered to have undergone range expansion, whereas if it was negative, it was considered to have experienced range contraction. Species pairs were then classified according to whether both species had undergone range expansion and range contraction. Out of 1602 pairs, both members of 607 pairs underwent range contraction while both members of 242 pairs experienced range expansions (Supporting information Table S1).

### Statistical analysis

All analyses were conducted using R (R Development Core team 2020). To test the hypotheses that behavioural interference influenced recent changes in sympatry or syntopy, we fit phylogenetic generalised linear mixed models (PLMMs) using an MCMC approach in the R package MCMCglmm (Hadfield 2010). Relatively non-informative priors that correspond to an inverse-Wishart distribution were used for random effects. Flat priors were used for fixed effects (Hadfield 2014). In addition to the fixed effects described above, the initial values of sympatry (i.e., sympatry in 1997-2000) and syntopy (in 1997-2000) were included to control for initial levels of coexistence because (a) species pairs that are already highly sympatric/syntopic are less likely to become even more so than those with low levels of sympatry/syntopy (Supporting information Fig. S4), and (b) in the case of syntopy, initial levels of syntopy is a confounding variable with a known relationship to interspecific territoriality (Drury et al. 2020). Random effects included species identity and a maximum clade credibility phylogeny (Jetz et al. 2012), specifying the nodes representing the most recent common ancestor of a pair (see further details in Losin et al. 2016, Drury et al. 2020). Each model was run for two million iterations with a burn-in of 20,000 and a thinning interval of 1000. We repeated each model fit four times and verified model convergence using Gelman-Rubin diagnostics (Gelman and Rubin 1992).

Models with interaction terms between interspecific territoriality and hybridisation (i.e., where species pairs that engage in both forms of behavioural interference are coded as ‘1’ in the interaction term) and between interspecific territoriality and non-hybridisation (i.e., where species pairs that defend interspecific territories but do not hybridize are coded as ‘1’ in the interaction term) were also fitted, as these combinations of behavioural interference may have differing impacts on coexistence (Grether et al. 2017, Cowen et al. 2020). If syntopy is a proxy for the similarity in habitat use between species, those pairs that are highly syntopic and therefore have similar requirements will respond more to changes in habitat availability than those with dissimilar requirements. To control for potential changes in habitat availability, we ran additional models in which both sympatry and syntopy datasets were subset to contain only pairs that shared the same habitat type (n = 871). Then, the habitat type of either species was included as a fixed effect to determine if changes in habitat have contributed to changes in sympatry and syntopy.

## RESULTS

### Syntopy

We found that interspecific territorial behaviour predicted changes in syntopy of North American passerines between 1997 and 2019 (Table 1, Fig. 2). Consistent with the hypothesis that forms of behavioural interference that cause habitat partitioning should facilitate coexistence, interspecifically territorial species pairs showed larger increases in syntopy than non-interspecifically territorial pairs (Table 1, Fig. 2). We also found that pairs occupying the same habitat types increased in syntopy more than those occupying different habitats (Table 1, Fig. 2). Further analyses restricted to species found in the same habitat still recovered an effect of interspecific territoriality (Supporting information Table S2), suggesting that our findings are robust to the potential confounding effects of changes in habitat suitability over this time period. We also found that pairs that defend different classes of intraspecific territories exhibit larger increases in syntopy than pairs that defend the same class of territory (Table 1, Fig. 2). When we restricted analyses to species pairs that defend the same type of territory and included terms for the type of intraspecific territoriality, thereby accounting for a potential confounding effect of intraspecific territory, we still recovered a strong effect of interspecific territoriality on Δ_syntopy_ (Supporting information Table S3).

**Table 1.** Predictors of Δ_syntopy_ from phylogenetic logistic mixed models (n = 1602 species pairs). The median coefficient estimates from the posterior distribution, as well as 95% credibility intervals and MCMC derived p-values are shown. Shaded rows indicate fixed effects with 95% credibility intervals that do not overlap 0. pMCMC values are from one chain (results are similar across all chains). The mean phylogenetic signal (λ) for this model was 0.101 (95% CI =0.002, 0.343).

|  | Median | 95% CI |  | pMCMC |  |
| --- | --- | --- | --- | --- | --- |
| Intercept | 0.046 | -0.319 | 0.415 | 0.773 |  |
| Interspecifically territorial | 0.335 | 0.116 | 0.552 | 0.001 | ** |
| Hybridising | -0.061 | -0.294 | 0.181 | 0.613 |  |
| Same intraspecific territory type | -0.138 | -0.270 | -0.007 | 0.045 | * |
| Patristic distance | 0.061 | -0.058 | 0.205 | 0.251 |  |
| Proportion shared axes | 0.005 | -0.051 | 0.060 | 0.848 |  |
| Both cavity nesters | 0.226 | -0.192 | 0.654 | 0.275 |  |
| Same habitat | 0.125 | 0.029 | 0.224 | 0.012 | * |
| Mass difference | 0.043 | -0.020 | 0.108 | 0.183 |  |
| Bill length difference | -0.014 | -0.077 | 0.048 | 0.658 |  |
| Both undergone range expansion | 0.010 | -0.129 | 0.152 | 0.891 |  |
| Both undergone range contraction | -0.014 | -0.127 | 0.100 | 0.817 |  |
| Syntopy 1997-2000 | -0.468 | -0.515 | -0.422 | <0.0005 | *** |

Significance codes: <0.05 \*, <0.01 \*\*, <0.001 \*\*\*

**Figure 2.**
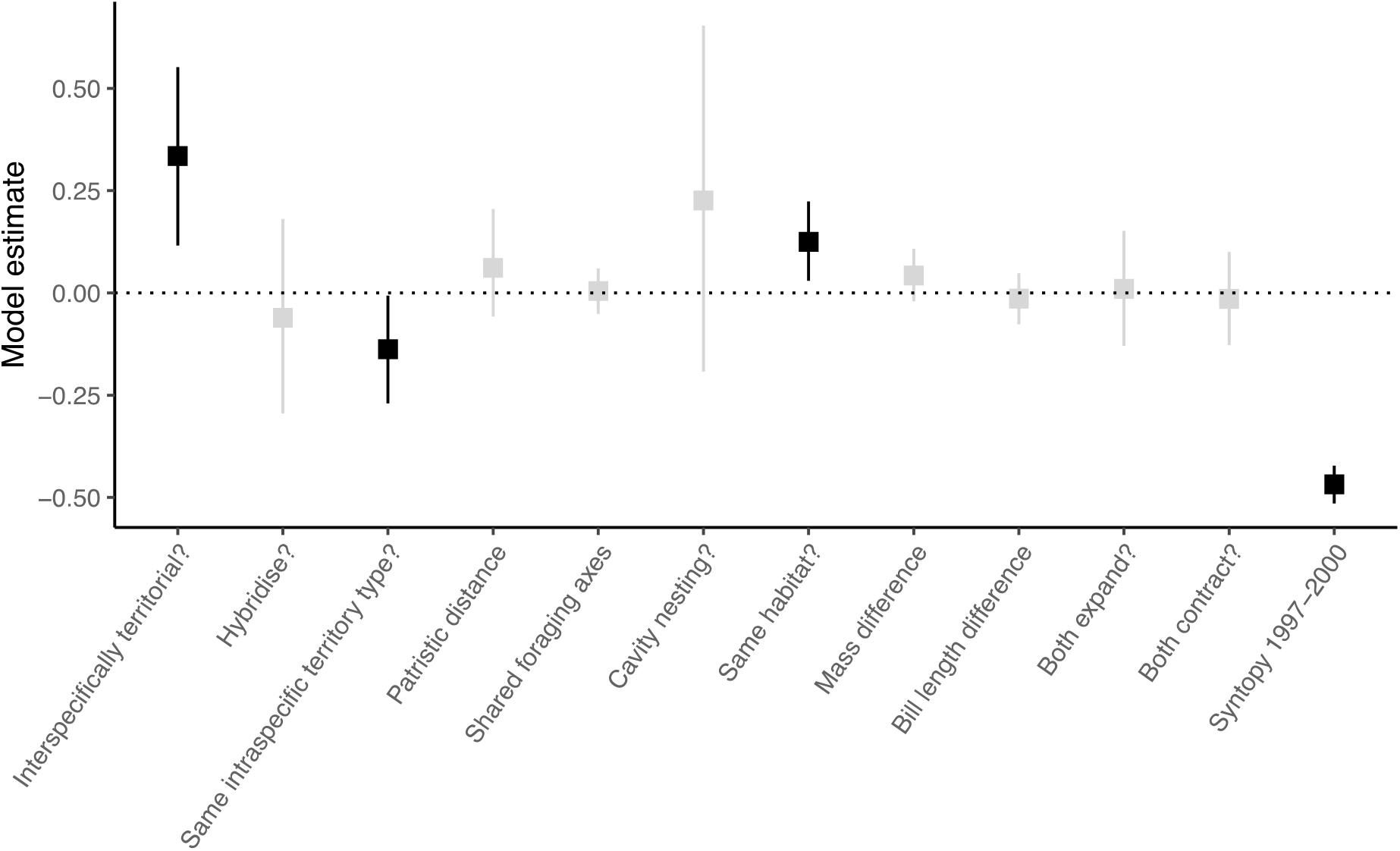
Coefficient estimates from a logistic regression (phylogenetic generalized linear mixed model) of Δ_syntopy_. Points correspond to the median and error bars represent the 95% credibility interval from four combined MCMC chains. Black points indicate fixed effects with estimates whose 95% credibility intervals do not include 0.

We found no effect of hybridization on Δ_syntopy_ (Table 1, Fig. 2), nor did we find any evidence that the interaction between interspecific territoriality and hybridization influenced syntopy (Supporting information Table S4).

### Sympatry

After controlling for range expansions and contractions, we found no evidence for an effect of behavioural interference on Δ_sympatry_ (Table 2, Fig. 3). Both mass difference and bill length difference were associated with changes in sympatry. As mass difference increased, Δ_sympatry_ decreased. Contrastingly, as bill length difference increased, Δ_sympatry_ increased. As with syntopy analyses, we found no effect of either hybridization or an interaction between interspecific territoriality and hybridization on Δ_sympatry_ (Table 2, Fig. 3, Supporting information Table S4).

**Table 2.** Predictors of Δ_sympatry_ from phylogenetic logistic mixed models (n = 1602 species pairs). The median coefficient estimates from the posterior distribution, as well as 95% credibility intervals and MCMC derived p-values are shown. Shaded rows indicate fixed effects with 95% credibility intervals that do not overlap 0. pMCMC values are from one chain (results are similar across all chains). The mean phylogenetic signal (λ) for this model was 0.025 (95% CI =0.0005, 0.162).

|  | Median | 95% CI |  | pMCMC |  |
| --- | --- | --- | --- | --- | --- |
| Intercept | 0.016 | -0.265 | 0.287 | 0.885 |  |
| Interspecifically territorial | 0.017 | -0.182 | 0.214 | 0.857 |  |
| Hybridising | 0.071 | -0.138 | 0.283 | 0.509 |  |
| Same intraspecific territory type | 0.052 | -0.099 | 0.200 | 0.494 |  |
| Patristic distance | -0.030 | -0.125 | 0.065 | 0.437 |  |
| Proportion shared axes | -0.013 | -0.071 | 0.044 | 0.660 |  |
| Both cavity nesters | 0.034 | -0.404 | 0.443 | 0.891 |  |
| Same habitat | -0.033 | -0.132 | 0.068 | 0.519 |  |
| Mass difference | -0.071 | -0.133 | -0.008 | 0.029 | * |
| Bill length difference | 0.069 | 0.011 | 0.128 | 0.022 | * |
| Both undergone range expansion | 0.425 | 0.250 | 0.595 | <0.0005 | *** |
| Both undergone range contraction | -0.247 | -0.399 | -0.097 | 0.001 | ** |
| Sympatry 1997-2000 | -0.102 | -0.147 | -0.056 | <0.0005 | *** |

Significance codes: <0.05 \*, <0.01 \*\*, <0.001 \*\*\*

**Figure 3:**
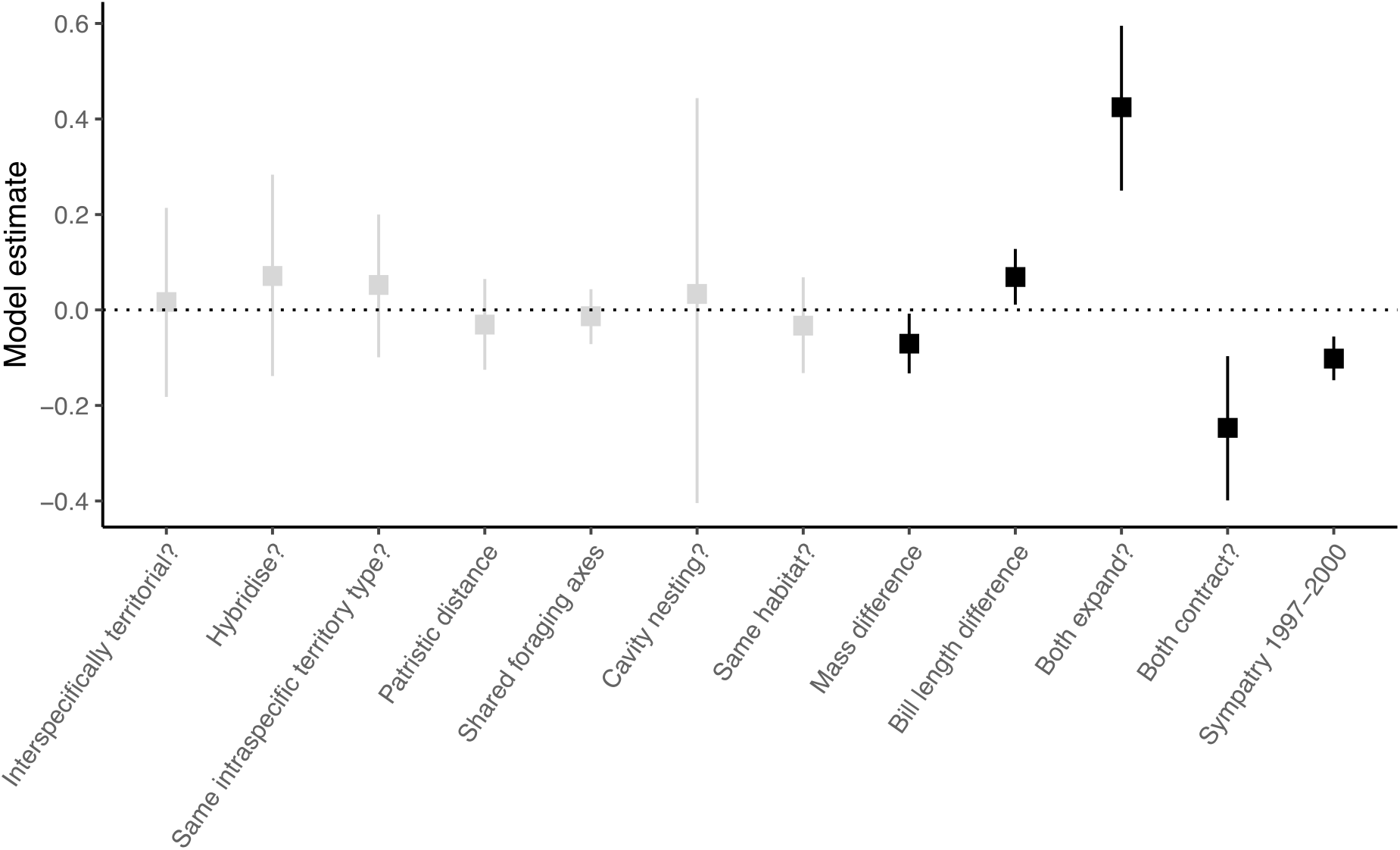
Coefficient estimates from a logistic regression (phylogenetic generalized linear mixed model) of Δ_sympatry_. Points correspond to the median and error bars represent the 95% credibility interval from four combined MCMC chains. Black points indicate fixed effects with estimates whose 95% credibility intervals do not include 0.

## DISCUSSION

We found support for the hypothesis that interspecific territoriality has accelerated fine-scale habitat overlap in North American passerines. To our knowledge, ours is the first study examining temporal changes in syntopy on a continental scale, showing a clear effect even over a short time period (22 years). The finding that interspecific territoriality facilitates fine-scale habitat overlap echoes several recent investigations demonstrating that behavioural interference affects spatiotemporal dynamics of assemblages (Gross and Price 2000, Duckworth and Badyaev 2007, Jankowski et al. 2010, Vallin et al. 2012, McQuillan and Rice 2015, Rybinski et al. 2016)

Other factors, in addition to behavioural interference, may have impacted the patterns of range dynamics that we documented. For instance, we found evidence that habitat type affected changes in syntopy, in line with previous studies finding that bird population dynamics in North America are not uniform across different habitat types (La Sorte and Boecklen 2005b, Rosenberg et al. 2019). Similarly, we found that defending the same intraspecific territory type was negatively associated with Δ_syntopy_. Given that the vast majority of North American passerines defend multipurpose territories, a non-zero value for this variable largely (in 97.5% of cases) represents cases where both species defend multipurpose territories, which may suggest that strong intraspecific territory defense on average slows down spatial movement (an important avenue for future research). Nevertheless, even after accounting for these factors, we found an effect of interspecific territoriality on recent shifts in fine-scale habitat overlap.

Models of sympatry dynamics, however, did not support a role for behavioural interference driving shifts in range-wide overlap. This finding supports the notion that simple indices of range overlap may fail to adequately capture the potential for species interactions (Drury et al. 2020). At a continental level, processes tend to be expressed on an evolutionary timescale rather than the ecological timescale of habitat-scale processes (Connor and Bowers 1987) and as such the relatively brief time span of this study may have failed to detect these processes. Nevertheless, we found evidence that traits related to resource competition are associated with changes in range-wide overlap; mass and bill similarity did predict changes in sympatry, though in opposing directions (mass similarity corresponded to increases in range overlap and bill length similarity corresponded to decreases in range overlap). The relationship between mass and range overlap could be related to higher dispersal ability in larger birds (Pigot and Tobias 2015) and/or habitat filtering (Polo and Carrascal 1999). The contrasting relationship between bill length and range overlap, on the other hand, could indicate a role for competitive exclusion (Pigot and Tobias 2013). Yet, why these effects would only act at a range-wide scale and not a local habitat scale is not clear. Further work could shed more light on whether these patterns hold at larger timescales or across other functional traits.

We found no evidence in models of syntopy or sympatry dynamics that either hybridization or the joint action of hybridization and interspecific territoriality impacted range dynamics. In contrast, recent analyses on sister-taxa suggest that behavioural interference, including hybridization, influences the attainment of secondary sympatry (Cowen et al. 2020). The discrepancy between our findings and those of Cowen et al. (2020) are likely a result of the vastly different timescales of our analyses. Indeed, most studies investigating the influence of species interactions on range dynamics concern changes that have taken place over millions of years (Gross and Price 2000, Jankowski et al. 2010, 2012, Pigot and Tobias 2013, 2015, Freeman et al. 2016, Boyce and Martin 2019). Nevertheless, some studies have focused on the influence of species interactions on contemporary range dynamics (Poling and Hayslette 2006, Duckworth and Badyaev 2007, Mac Nally et al. 2012, van Dongen et al. 2013, Wiens et al. 2014, Friedemann et al. 2017). Another consideration that may explain discrepancies between ours and other studies relates to starting levels of interspecific range overlap. Theoretical models posit that range dynamics in the face of behavioural interference are likely to be positively frequency-dependent, and therefore, systems with interference are often prone to competitive and/or sexual exclusion when at least one species occurs at a low frequency (Kuno et al. 1992). The species pairs studied here, however, were already sympatric at the onset of the study period, and therefore interspecific territoriality between these lineages is likely to be evolutionarily stable (Drury et al. 2020, Cowen et al. 2020).

Determining how processes that unfold on ecological timescales scale up to generate macroevolutionary dynamics is an open challenge in the field (Weber et al. 2017, Harmon et al. 2019, Hembry and Weber 2020). Further research harnessing the power of long-term census data like those from the North American Breeding Bird Survey promises to play an important role in achieving this micro-to-macro link. Our analyses contribute to a growing body of work demonstrating how behavioural interference impacts fundamental ecological and evolutionary processes. Further understanding of these impacts will improve our ability to predict the consequences of species interactions that form in novel assemblages as a result of ongoing, human-induced global change.

## Data archiving

The data files and scripts used in analyses will be uploaded to Dryad upon initial acceptance.

## Acknowledgements

We thank Christophe Patterson for helpful comments on the manuscript.

