## Supporting Information for "Interspecific territoriality has facilitated recent increases in the breeding habitat overlap of North American passerines"

Tables S1-S4
Figures S1-S4

**Table S1.** The number of pairs in which the constituent species expanded, contracted, or had no net change in the number of BBS routes between 1997 and 2019.

| | | Number of pairs | | |
|---|---|---|---|---|
| | | Species 1 | | |
| | | Range expansion | Range contraction | No change |
| Species 2 | Range expansion | 242 | 408 | 14 |
| | Range contraction | 271 | 607 | 21 |
| | No change | 16 | 21 | 2 |

**Table S2.** Predictors of $\Delta_{syntopy}$ when restricted to pairs that share the same habitat type (n = 871 species pairs). The median coefficient estimates from the posterior distribution, as well as 95% credibility intervals and MCMC derived p-values are shown. Shaded rows indicate fixed effects with 95% credibility intervals that do not overlap 0. pMCMC values are from one chain (results are similar across all chains). The mean phylogenetic signal (λ) for this model was 0.075 (95% CI = 0.001, 0.347).

| | Median | 95% CI | | pMCMC | |
|---|---|---|---|---|---|
| Intercept | 0.176 | -0.233 | 0.598 | 0.331 | |
| Interspecifically territorial | 0.263 | 0.012 | 0.521 | 0.044 | * |
| Hybridising | 0.080 | -0.200 | 0.369 | 0.558 | |
| Same intraspecific territory type | 0.027 | -0.159 | 0.217 | 0.750 | |
| Patristic distance | 0.068 | -0.049 | 0.226 | 0.226 | |
| Proportion shared axes | -0.055 | -0.128 | 0.017 | 0.137 | |
| Both cavity nesters | 0.046 | -0.018 | 0.111 | 0.142 | |
| Intermediate habitat | -0.153 | -0.450 | 0.141 | 0.302 | |
| Complex habitat | -0.241 | -0.516 | 0.031 | 0.075 | |
| Mass difference | 0.042 | -0.048 | 0.130 | 0.350 | |
| Bill difference | 0.014 | -0.067 | 0.097 | 0.757 | |
| Both undergone range expansion | 0.007 | -0.172 | 0.190 | 0.971 | |
| Both undergone range contraction | -0.040 | -0.188 | 0.106 | 0.568 | |
| Syntopy 1997-2000 | -0.468 | -0.531 | -0.405 | <0.0005 | *** |

Significance codes: <0.05 *, <0.01 **, <0.001 ***

**Table S3.** Predictors of $\Delta_{syntopy}$ when restricted to pairs that share the same intraspecific territory type (n = 1221 species pairs). The median coefficient estimates from the posterior distribution, as well as 95% credibility intervals and MCMC derived p-values are shown. Shaded rows indicate fixed effects with 95% credibility intervals that do not overlap 0. pMCMC values are from one chain (results are similar across all chains). The mean phylogenetic signal (λ) for this model was 0.239 (95% CI = 0.012, 0.643).

| | Median | 95% CI | | pMCMC | |
|---|---|---|---|---|---|
| Intercept | -2.196 | -3.356 | -1.238 | <0.0005 | *** |
| Interspecifically territorial | 0.329 | 0.088 | 0.567 | 0.007 | ** |
| Hybridising | -0.037 | -0.290 | 0.212 | 0.802 | |
| Intraspecific territoriality 4 | 2.036 | 1.136 | 3.042 | <0.0005 | *** |
| Intraspecific territoriality 5 | 2.146 | 1.290 | 3.138 | <0.0005 | *** |
| Patristic distance | 0.070 | -0.161 | 0.276 | 0.419 | |
| Proportion shared axes | 0.041 | -0.023 | 0.104 | 0.202 | |
| Both cavity nesters | -0.035 | -0.501 | 0.435 | 0.886 | |
| Same habitat | 0.154 | 0.040 | 0.267 | 0.011 | * |
| Mass difference | 0.047 | -0.024 | 0.116 | 0.185 | |
| Bill difference | 0.000 | -0.066 | 0.069 | 0.998 | |
| Both undergone range expansion | 0.067 | -0.090 | 0.225 | 0.380 | |
| Both undergone range contraction | 0.083 | -0.049 | 0.207 | 0.201 | |
| Syntopy 1997-2000 | -0.454 | -0.506 | -0.401 | <0.0005 | *** |

Significance codes: <0.05 *, <0.01 **, <0.001 ***

**Table S4.** Model selection on analyses that include all species pairs does not support any interaction between interspecific territoriality and hybridisation. Fixed effects are interspecific territoriality, hybridisation, same intraspecific territory type, patristic distance, proportion of shared foraging axes, both cavity nesters, same habitat, square root mass difference, square root bill length difference, both undergone range expansion, both undergone range contraction and mean syntopy in 1997-2000. In each model, there were 1602 species pairs, of which 74 were interspecifically territorial. We included different models for two possible interactions between interspecific territoriality and hybridisation, as these combinations of behavioural interference may have contrasting impacts on coexistence (see details in main text).

| | DIC mean (range) | |
|---|---|---|
| Model terms | Syntopy | Sympatry |
| Fixed effects (no interactions) | 4071.20 (4071.09-4071.29) | 3732.59 (3732.53-3732.65) |
| Fixed effects + interspecifically territorial*hybridising | 4071.24 (4071.21-4071.27) | 3732.16 (3732.12-3732.20) |
| Fixed effects + interspecifically territorial*non-hybridising | 4071.23 (4071.17-4071.33) | 3732.15 (3732.07-3732.22) |

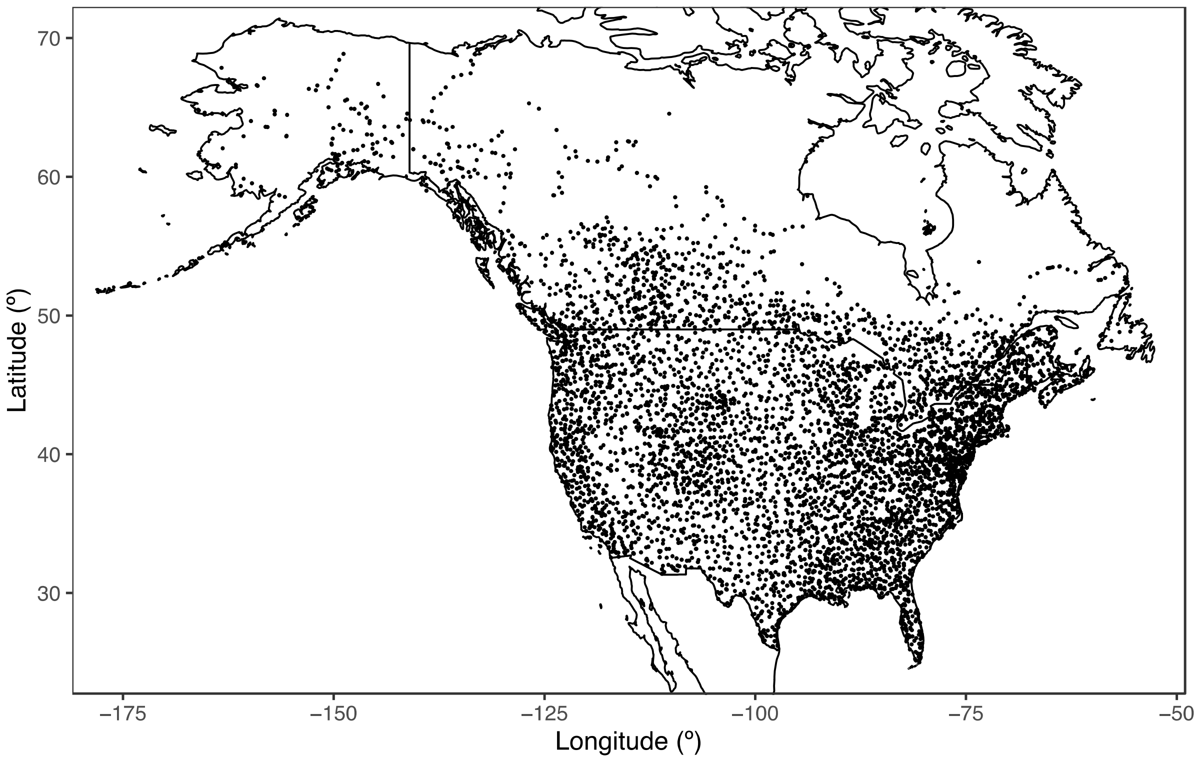


**Figure S1**. Map of all North American Breeding Bird Survey routes between 1966 and 2019 (n=5756), not every route has been surveyed annually.

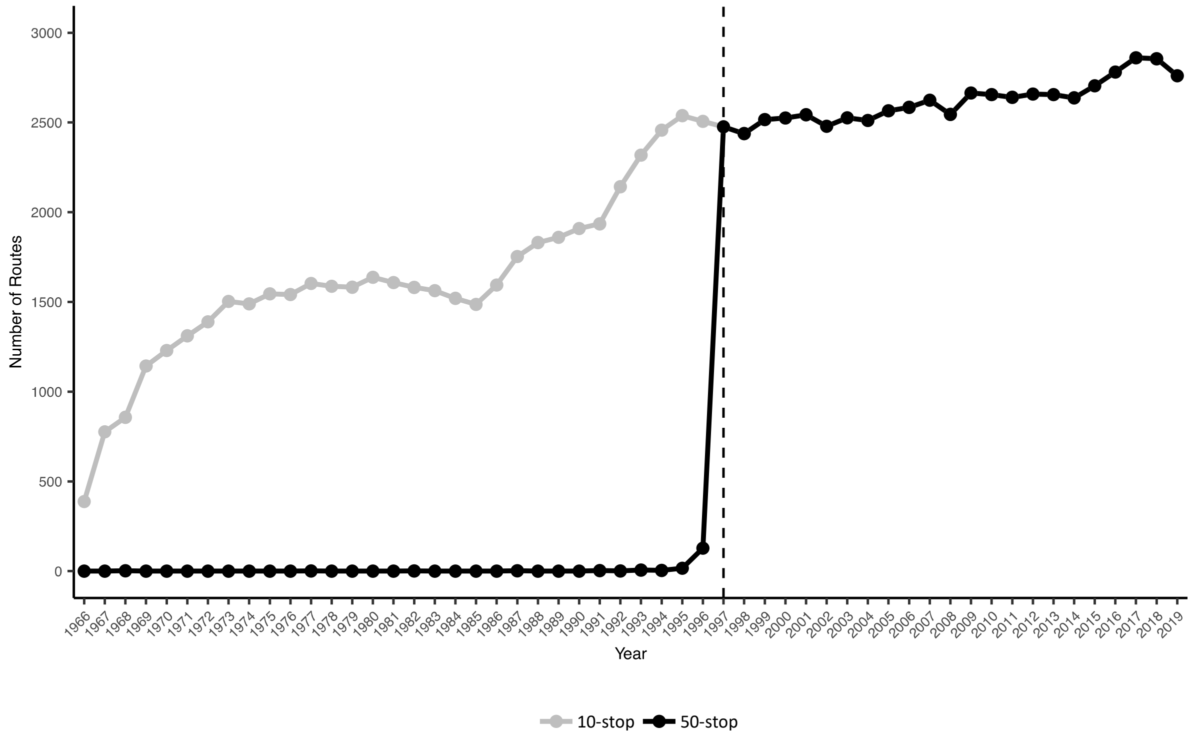


**Figure S2**. Number of routes for which there is 10-stop (grey), used for sympatry analyses, and 50-stop (black) data, necessary for syntopy analyses, available for each year between 1966 and 2019.

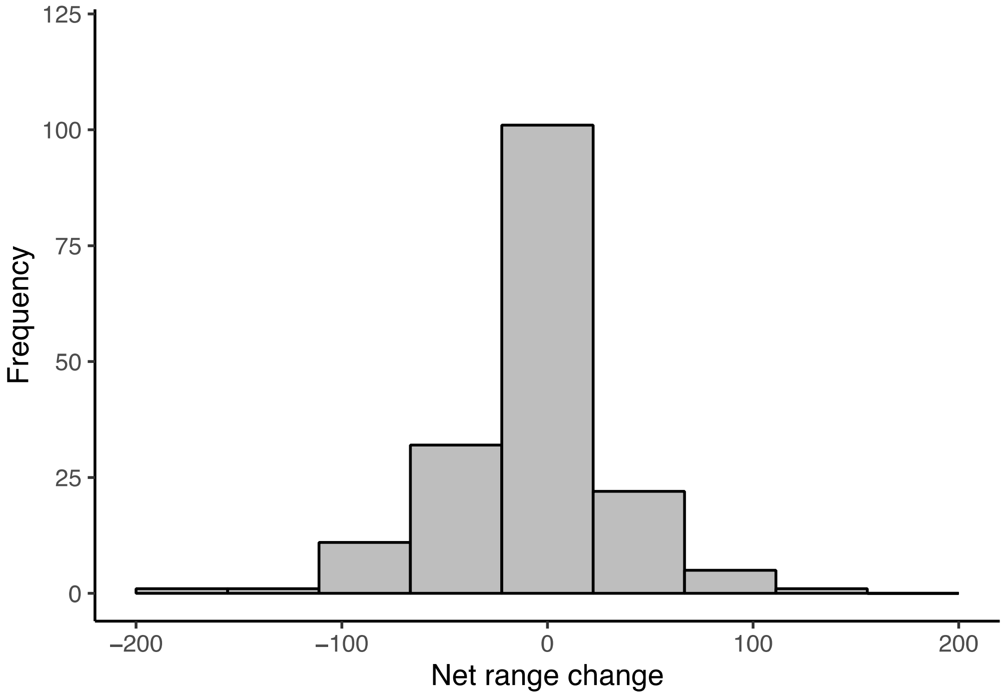


**Figure S3**: Net range change of 174 species. Net range change is calculated as the difference between the number of route gains minus the number of route losses for a given species between 1997 and 2019

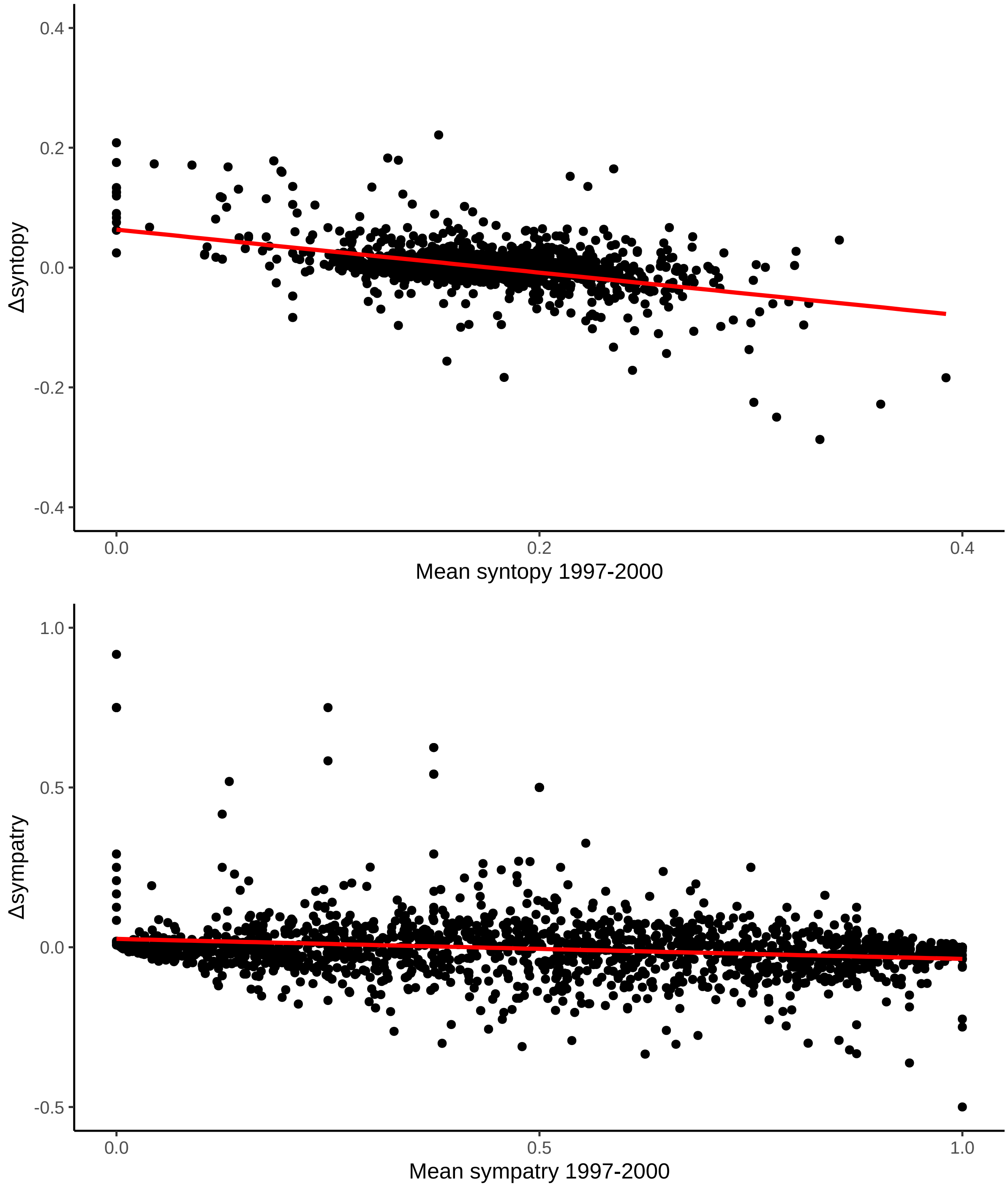


**Figure S4.** Relationship between Δsyntopy and mean syntopy between 1997-2000 (top) and relationship between Δsympatry and mean sympatry between 1997-2000 (bottom).
